# Mechanosensitive binding of p120-Catenin at cell junctions regulates E-Cadherin turnover and epithelial viscoelasticity

**DOI:** 10.1101/357186

**Authors:** K. Venkatesan Iyer, Romina Piscitello-Gómez, Frank Jülicher, Suzanne Eaton

## Abstract

Studying how epithelia respond to mechanical stresses is key to understanding tissue shape changes during morphogenesis. Here, we study the viscoelastic deformation of the Drosophila pupal wing epithelium in response to mechanical stress that evolves during morphogenesis. We show that wing epithelial tissue viscoelasticity depends on endocytic turnover of E-Cadherin. The fraction of ECadherin undergoing turnover depends on mechanical stress in the epithelium. We identified mechanosensitive binding of the endocytic regulator p120-Catenin (p120) as a mechanism to regulate E-Cadherin turnover. Under high stress, p120 is released into the cytoplasm, destabilizing E-Cadherin complexes and increasing its turnover. In p120 mutants, E-Cadherin turnover is insensitive to mechanical stress. Furthermore, we show that p120 is crucial for the viscoelastic deformation of the wing epithelium. Taken together, our findings reveal that mechanosensitive binding of p120-Catenin tunes epithelial tissue viscoelasticity during morphogenesis.

## Introduction

How tissue size and shape emerge from the collective behaviour of cells is a key question in developmental biology. Time lapse imaging have revealed that developing tissues reorganize dynamically, indicating fluid like aspects in their material properties (Blanchard et al., 2009; Aigouy et al., 2010; Etournay et al., 2015; Jülicher and Eaton, 2017). Tissue shape emerges from a dynamic interplay between patterns of active mechanical stress generation, and the response of cells and tissues to mechanical stresses (Lecuit et al., 2011; Guirao and Bellaïche, 2017). Most studies have focussed on how tissue dynamics results from patterns of cellular force generation organized by biochemical signals (Heisenberg and Bellaïche, 2013; Naganathan et al., 2014; Munjal et al., 2015). However, the possible mechanical responses of tissues, (Chanet and Martin, 2014), and how these responses are deployed during development, remain largely unclear.

The response of soft materials, like tissues to mechanical stress depends on their material properties, which are typically viscoelastic (Cross, 2012; Forgacs et al., 1998; Kumar et al., 2006). Elastic materials deform in response to mechanical stress and return to their original shape when stress is removed. Viscous materials behave like fluids – they relax mechanical stresses by remodeling their internal structure and do not return to their original shape after deformation. Viscoelastic materials combine both behaviors– they are elastic over short timescales and viscous at longer timescales. What mechanisms determine the viscoelastic properties of developing tissues? Studying such mechanisms is key to understanding stress-dependent tissue shape changes during development.

The *Drosophila* pupal wing is an ideal system to study how epithelial tissues respond to mechanical stress during morphogenesis (Aigouy et al., 2010; Etournay et al., 2015). From 16 hours after puparium formation (hAPF), the wing blade epithelium deforms in response to anisotropic proximal-distal (PD)-oriented tissue stresses. These stresses develop during contraction of the more proximal hinge region. Stresses develop because the wing blade is connected along its margin to an overlying cuticle through an extracellular matrix protein called Dumpy (Etournay et al., 2015; Ray et al., 2015; Wilkin et al., 2000). Over about 20 hours, the wing blade extends along the PD axis and narrows in the anterior-posterior (AP) axis. Decomposing the overall tissue deformation into contributions stemming from different cellular processes revealed that oriented cell divisions, cell rearrangements and cell shape changes all contribute to this tissue shape change (Etournay et al., 2015, 2016; Merkel et al., 2017). One key missing piece of information is the extent to which the combination of cell mechanics and cell rearrangements relax the stresses generated by hinge contraction. Furthermore, the molecular mechanisms that underlie stress-dependent changes in cell shape and cell rearrangements are not known.

Cell shape changes and rearrangements during morphogenesis involve the remodeling of the apical junctional network (Baum and Georgiou, 2011; Takeichi, 2014). E-Cadherin is a core component of *adherens* junctions that not only mediates adhesion between cells, but also regulates the linkage of adhesive complexes to the underlying acto-myosin cytoskeleton and controls cytoskeletal dynamics (Lecuit and Yap, 2015; Takeichi, 2014). Most studies in cultured cells have highlighted how mechanical stress increases the linkage of E-Cadherin to the actin cytoskeleton and promotes adhesion. a-Catenin, which is recruited to the E-Cadherin cytoplasmic tail through b-catenin, unfolds under stress and binds to actin filaments (Buckley et al., 2014). a-Catenin also recruits another mechanosensitive protein, Vinculin, which upon unfolding enhances actin binding and actin polymerization (Seddiki et al., 2017; Yao et al., 2014). However, results in MDCK cells suggest that stress can also destabilize E-Cadherin dependent adhesions (Beco et al., 2015). This appears to involve internalization of E-Cadherin complexes, but the mechanisms underlying stress dependent internalization are not known. Such mechanisms could be particularly relevant during morphogenesis of the pupal wing, where stress induces remodeling of the junctional network. Therefore, they may also be important for the understanding of viscoelastic behaviors of such tissues.

Here we show that the *Drosophila* wing blade exhibits viscoelastic behaviors upon deformation in response to stress induced by hinge contraction. Both cell rearrangements and changes in cell mechanics contribute to the viscosity of the wing epithelium. We provide evidence that mechanical stresses destabilize E-Cadherin complexes and elevate endocytic turnover of E-Cadherin in the wing blade. Mechanosensitivity of E-Cadherin turnover depends on p120-Catenin (p120), a protein that binds to the E-Cadherin tail and blocks access to the endocytic machinery. We show that p120 is released from *adherens* junctions when stress is high, and re-associates with junctions as stress relaxes. In the p120 mutant, E-Cadherin turnover no longer responds to tissue stress and remains high even when stresses relax. The apparent viscosity of the wing epithelium is reduced by loss of p120. Our work reveals that mechanosensitive binding of p120 regulates the turnover of E-Cadherin and thereby tunes the viscoelastic properties of the wing epithelium.

## Results

### Epithelial tissue undergoes viscoelastic deformation during morphogenesis

The first step towards studying the viscoelastic properties of the wing is to measure the time dependence of mechanical stress in the wing during morphogenesis. Focusing on the region between the 3^rd^ and 4^th^longitudinal veins, we used two laser-ablation approaches to estimate the orientation and magnitude of mechanical stress at different times of development (Figure 1A). To reveal stress orientation, we performed circular laser ablations and monitored the deformation of the cut shape over time (Figure 1B). Circular cuts deformed into ellipses whose major axes were predominantly aligned with the PD axis of the wing, confirming that mechanical stress remains aligned with the PD axis throughout morphogenesis (Figures 1C and 1D). These experiments also suggested that the magnitude of mechanical stress first increases and then decreases over time (Figure S1). The initial velocity of retraction is the best proxy for stress (Mayer et al., 2010). However, because of the time required to perform circular cuts, it is difficult to measure this velocity for such cuts. We therefore performed linear ablations orthogonal to the PD axis to measure the initial velocity of retraction at different times (Figure 1E). These studies revealed that stress peaks at 20 hAPF and then gradually relaxes back to its initial low value by 32 hAPF (Figure 1F). The fact that stress relaxes during wing morphogenesis indicates that the wing blade has viscous properties. Our analysis further suggests that not only cell rearrangements, but changes in preferred cell shape – the shape of cells in a tissue not under mechanical stress, also contribute to stress relaxation during morphogenesis (Figures 1G and 1H).

**Figure 1:**
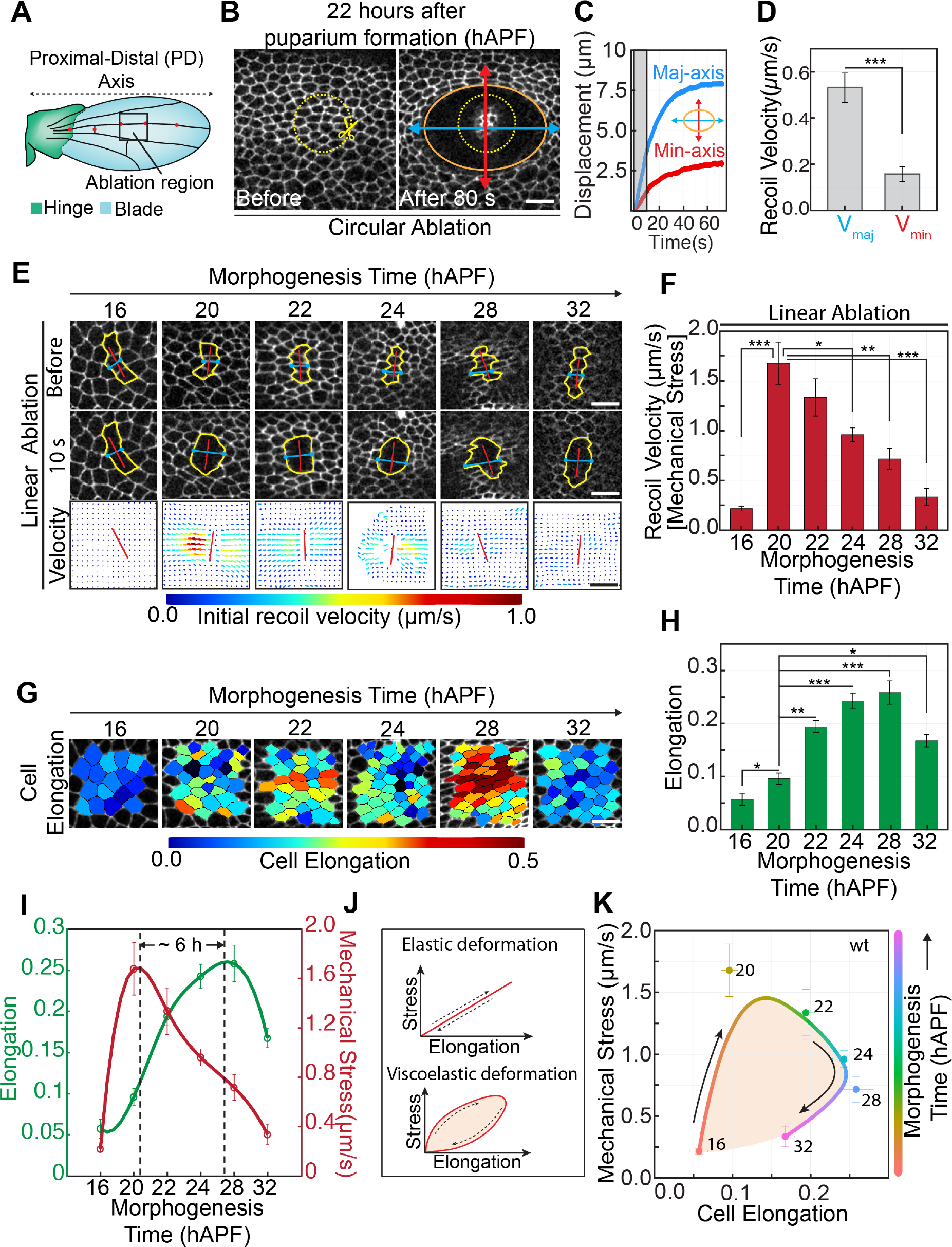
Epithelial tissue undergoes viscoelastic deformation during morphogenesis. **(A)** Schematic of the Drosophila pupal wing epithelium showing the hinge (green) and blade (blue) regions. Red dots represent sensory organs. Laser ablations were performed in the region shown in black ROI. **(B)** Images of the region of the pupal wing shown in (A), before and 80 s after circular ablation of 20 µm diameter at 22 hAPF. Yellow dotted ROI (with scissors) represents the ablated circle. Orange ellipse shows the size of the ablated region 80 s after ablation. Blue and red double-sided arrows show the expansion along major and minor axes of the ellipse. Scale bar, 10 µm. **(C)** Displacement vs time graph for the circular ablation. Blue and red curves represent displacement along the major and minor axis of the ellipse respectively. Grey region represents the region of the curve used for estimating the initial recoil velocity. **(D)** Initial recoil velocities along the major and minor axes of the ellipse. Error bars, SEM. *** represents p < 0.0001. (n≥ 5 wings) **(E)** Images showing linear laser ablation in the region of pupal wing shown in (A), during morphogenesis. Red dotted line represents the 7 µm ablation. Yellow ROI represents the outline of the cells around the ablation line, before and 10 s after ablation. Velocity image shows the recoil velocity profile with velocity vectors originating from the site of ablation. The length and color of the vectors represents the magnitude of the recoil velocity. Scale bar, 10 µm. **(F)** Bar graph shows the recoil velocity (proxy for mechanical stress) during different stages of the wing morphogenesis. Error bars, SEM. * p< 0.05, ** p<0.001, and *** p< 0.0001. P-values were estimated with respect to 20hAPF. (n≥ 5 wings in each case) **(G)** Color coded images showing cell elongation at different stages of wing morphogenesis for the same region as laser ablation. Scale bar, 5 µm. **(H)** Bar graph shows the cell elongation during wing morphogenesis. Error bars, SEM. * p< 0.05, ** p<0.001, and *** p< 0.0001. P-values were estimated with respect to 20hAPF. (n ≥5wings in each case). **(I)** Cell elongation (green) and mechanical stress (dark red) during wing morphogenesis. Open circles show the experimental data and the line represents spline fit to the data. The peak of mechanical stress and cell elongation are separated by a 6 hr time delay. Error bars, SEM **(J)** Schematic showing the relationship between stress and elongation during cyclic stretching of an elastic and viscoelastic material. **(K)** Mechanical stress vs cell elongation graph during pupal wing morphogenesis. Filled circles represent experimental data, color coded for the time of morphogenesis. Numbers beside the points show the stage of wing morphogenesis (in hAPF). The smooth curve represents the spline fit to the data color coded for morphogenesis time. The area under the smooth curve (peach) shows the hysteresis in the process and corresponds to the dissipation due to viscoelastic deformation. Scale bar, 10µm. Error bars represent S.E.M. See also Figure S1.

We took advantage of the increase and decline of tissue stress during wing tissue deformation to investigate tissue viscoelastic properties. Viscoelasticity of soft materials is typically characterized by studying the relationship between periodic material deformations and corresponding oscillating stress. An elastic material undergoes instantaneous deformation that is in phase with the applied stress oscillation. A viscous material shows a time delay between the deformation and applied stress such that deformation and stress are 90 degrees phase shifted. Viscoelastic materials show a response with an intermediate phase shift. Measuring deformation during mechanical loading and unloading permits one to plot a mechanical stress-deformation curve which provides information about the phase relation between deformation and stress, and thus the viscoelasticity of the tissue (Figure 1J). To study the relationship between cell deformation and mechanical stress, we measured the average cell elongation in the region around the ablation prior to the ablation. As previously described, cells elongate in the PD axis during the first phase of morphogenesis, and then partially relax their shape as cell rearrangements help to relax stress and contribute to PD axis elongation (Figures 1G and 1H). Plotting both average cell elongation and mechanical stress as a function of time revealed a delay of 6 hours between the peaks of stress and cell deformation (Figure 1I), suggesting that cells themselves exhibit viscoelastic behaviors. The viscous component is also highlighted by the large area enclosed by the stress-deformation curve (Figure 1K). These results show that changes in preferred cell shape in the pupal wing contribute to viscoelastic stress relaxation.

## E-Cadherin turnover during morphogenesis is mechanosensitive

To begin the investigation of molecular mechanisms that underlie the viscoelastic properties of the wing epithelium, we examined the rate of E-Cadherin turnover during morphogenesis. To do so, we used E-Cadherin GFP to perform fluorescence recovery after photobleaching (FRAP) measurements at different times during pupal morphogenesis (Figure 2A). We photobleached a single cell junction in the same region and at the same times at which we had measured mechanical stress using laser ablation, and then quantified the recovery of fluorescence over a period of 500 seconds (Figures 2B, 2C and Figure S2). Recovery over this time period was well described by a double exponential function (Table S1),

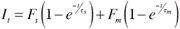

where *I_t_* is the intensity of the junction at time *t*, *τ_s_* and *τ_m_* are two timescales of recovery, and *Fs* and *F_m_* are the corresponding recovery fractions. The timescale *τ_s_* ranged from 10-50 seconds and is referred to as the second-timescale (Figure S3B). *F_s_* is the corresponding second-scale recovery fraction. The timescale *τ_m_* varied between 2-8 minutes, and is referred to as the minute-timescale (Figure S3A). *F_m_* is the corresponding minute-scale recovery fraction. In longer movies, an additional recovery process was observed, revealing a third timescale on the order of ˜ 2 hours (Figure S4). We refer to this fraction as the immobile fraction. As the mechanical stress increases during 16-22 hAPF, *F_m_* is consistently high and increases slightly, peaking at 22 hAPF. As mechanical stress drops between 22 and 32 hAPF, *F_m_* also decreases, while the immobile fraction increases (Figure 2D and Figure S5). We observed a significant correlation between mechanical stress and *F_m_* (Figure 2E). In contrast, *F_s_* does not vary strongly during this process. These observations suggest that minute-scale recovery fraction and immobile fraction are mechanosensitive, whereas the second-scale fraction is insensitive to mechanical stress.

**Figure 2:**
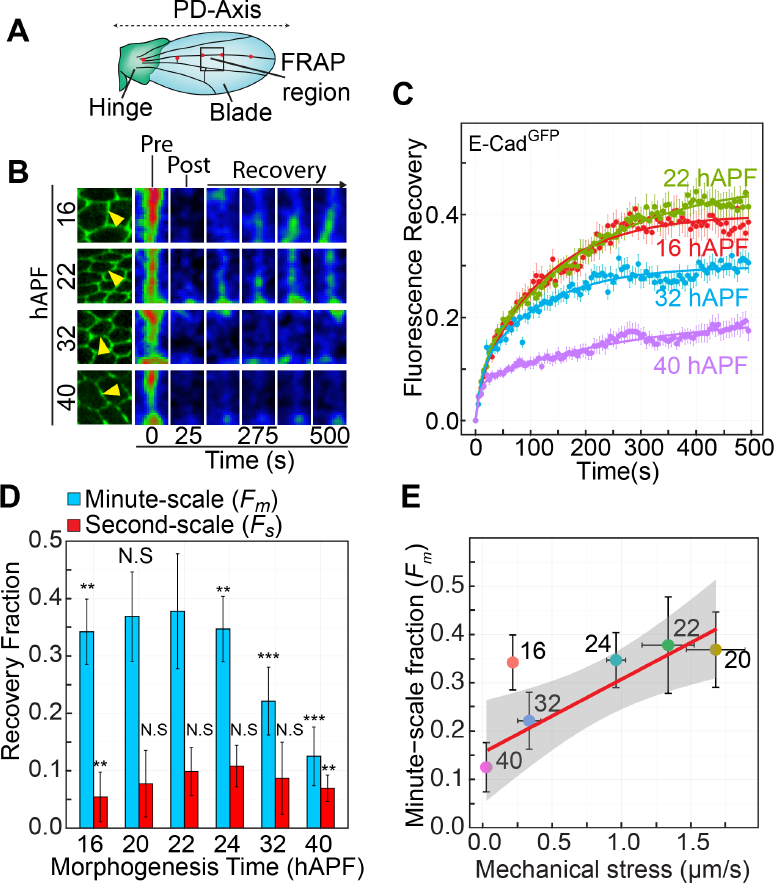
E-Cadherin turnover during morphogenesis is mechanosensitive. **(A)** Schematic shows that FRAP experiments were performed in the same region of the pupal wing where laser ablations were performed. **(B)** Time-lapse images from FRAP time series of E-Cad^GFP^ at 16, 22, 32 and 40hAPF. Yellow arrows show the junctions that are bleached. Time series images show the pre- and post-bleach images and the recovery over 500 s. Images are color coded for intensity. **(C)** FRAP curves for E-Cad. Solid line shows the double exponential fit. Error bars, SEM.(n≥ 40 junctions in each case) **(D)** Minute-scale, *F_m_* (blue) and second-scale, *F_s_* (red) recovery fractions for E-Cad. Error bars, S.E.M. ** p < 0.01 and *** p < 0.001. P-values in are estimated for *F_m_* and *F_s_* at all time points with respect to 22 hAPF. N.S indicates that the differences are not significant. (n≥ 40 junctions in each case) **(E)** Scatter plot between mechanical stress and minute-scale recovery fraction (*F_m_*). Color of the points and the numbers beside the points represent time of morphogenesis in hAPF. Solid black line shows the linear fit with 16 hAPF excluded and the grey shaded region represents the error in the fit. See also Figures S2, S3, S4 and S5, and Table S1.

## Mechanical stress regulates E-Cadherin turnover

We have shown that change in mechanical stress during pupal wing morphogenesis correlates strongly with changes E-Cad turnover. One explanation for this correlation is that mechanical stress directly regulates this process. Alternatively, mechanical stress and E-Cadherin turnover might be controlled independently during morphogenesis. To resolve these possibilities, we used both genetic and physical means to reduce mechanical stress at a time when it is normally high, and then examined E-Cad turnover. First, we used the hypomorphic Dumpy mutant, *dp^ov1^*(Wilkin et al., 2000). In this mutant, matrix connections between the cuticle and the distal tip of the wing are depleted, and these wings therefore never develop the same magnitude of mechanical stress as wild type wings (Figure S6). This leads to reduced cell area and cell elongation (Figure 3A and Figure S6). We also depleted the levels of Dumpy in the entire wing by expressing dsRNA of Dumpy in the wing – *dp*^RNAi^ (Ray et al., 2015). This produces stronger adult phenotypes and affects cell elongation even more dramatically than *dp^ov1^* (Figure 3A Figure S6). We performed FRAP on single junctions of wild-type, *dp^ov1^* and *dp*^RNAi^ wings at 24 hAPF when Dumpy mutant wing blades have already retracted from the cuticle (Figures 3B and 3C). Both Dumpy perturbations strongly reduces E-Cad turnover, specifically the minute-scale recovery fraction, *F_m_* of E-Cad, while the second-scale fraction, *F_s_* is not affected (Figure 3D and Figure S7). These FRAP experiments suggest that mechanical stress increases the turnover of E-Cad.

**Figure 3:**
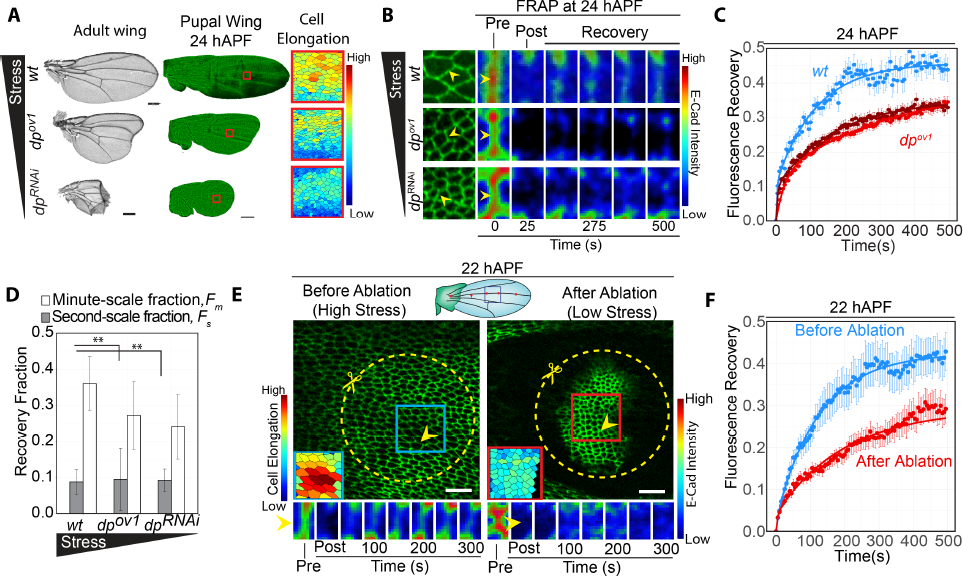
Mechanical stress regulates E-Cad turnover. **(A)** Adult wings (left panel) and 24 hAPF pupal wings (middle panel) of *wt, dp^ov1^* and *dp^RNAi^*.Scale bar in left panel, 200 µm. Scale bar in middle panel, 100µm. Red ROI represents the region for which color-coded cell elongation is shown in the right panel. **(B)** Images show a small region of the wing where FRAP was performed. Yellow arrows show the bleached junctions. Time lapse images show the pre- and post- bleach frames, followed by recovery of fluorescence. **(C)** Fluorescence recovery curves for *wt* (blue), *dp^ov1^* (red) and *dp^RNAi^* (dark red) at 24 hAPF. Solid lines represent double exponential fits. Error bars, SEM. (n≥ 65 junctions in each case). **(D)** Minute-scale (white) and second-scale (grey) recovery fractions for *wt, dp^ov1^* and *dp^RNAi^*.Error bars, S.E.M. ** p < 0.001. (n≥ 65 junctions in each case). **(E)** The schematic of the pupal wing shows the region where ablation was performed (black ROI in the wing). Yellow dotted ROI (with scissors) show the ablated region. Bottom insets show color coded cell area before ablation (blue ROI) and after ablation (red ROI). Scale bar, 10 µm. Bottom panels show the intensity color coded time lapse images of fluorescence recovery of the bleached junctions shown by yellow arrow, before and after ablation. **(F)** Fluorescence recovery curves before (blue) and after (red) ablation. Error bars, SEM. (n≥15 junctions in each case). See also Figures S6, S7, S8 and S9.

To investigate the influence of acute reduction in mechanical stress on E-Cad turnover, we performed FRAP experiments in wild type wings before and after a circular laser ablation at 22 hAPF, when the blade is under high mechanical stress (Figure 3E and Figure S8A). After the first FRAP measurements, we ablated a circle with a radius of 20 µm around the analyzed region. This isolated a small tissue island from the surrounding tissue (Figure 3E). Within 10 minutes of ablation, cells in the tissue island contracted their apical cross-section and became less elongated (Figure 3E), indicating that the laser ablation rapidly relieves mechanical stress in the tissue island. We then performed a second FRAP analysis within this isolated region. After ablation, *F_m_* decreased by 47±5% (Figure 3F and Figure S8B). In contrast, *F_s_* was not affected. Thus, while endocytic turnover of E-Cad responds rapidly to changes in stress, its diffusion within the membrane does not. We observed that E-Cad turnover was consistently high in early stage (16 hAPF) and did not increase further with increasing mechanical stress (Figures 2B, 2C and 2D). Thus, we wondered to what extent mechanical stress-dependent mechanisms contributed to E-Cad turnover at this stage. To investigate this, we measured E-Cad turnover at 16 hAPF before and after generating an isolated tissue island by circular laser ablation (Figures S8C and S8D). In contrast to 22 hAPF, only 26±5% of *F_m_* was stress dependent (Figures S8C, S8D and S8E). This revealed that at 16 hAPF, a significant but smaller fraction of *F_m_* is dependent on mechanical stress. However additional active mechanisms could also promote E-Cad turnover at this stage. Taken together, these data show that the mechanical stress regulates the turnover of E-Cad during wing morphogenesis.

We wondered whether stress-dependent turnover is specific to *adherens* junctions, or whether stress might also act at septate junctions to increase protein turnover. Septate junctions are located approximately 3 µm below the *adherens* junctions (Figure S9A) and are thought to inhibit the flux of small molecules and ions between epithelial cells (Banerjee et al., 2006). We performed laser ablations at the level of septate junctions, visualized using Neuroglian::EGFP (Nrg) (Figure S9B and S9C). Interestingly, the septate junction region is under significantly less mechanical stress than the *adherens* junction region containing E-Cad (Figure S9C). Furthermore, FRAP analysis indicated that Nrg turns over much more slowly than E-Cad and that turnover does not change during pupal wing remodelling (Figure S9D and S9E). Thus, E-Cad and the *adherens* junction region appear to be specialized sites of mechanical stress sensing and response.

## Membrane trafficking contributes to the minute-scale recovery of E-Cadherin

Next, we asked what cell biological mechanisms contribute to the minute-scale and second-scale recovery fractions. E-Cad fluorescence could in principle recover either through the vesicular delivery of new E-Cad from the cytoplasm or by lateral diffusion within the membrane (Bulgakova et al., 2013). To distinguish the relative contributions of each process, we used two different approaches to perform FRAP at 22 hAPF. In the first approach, we bleached the junction of interest (JOI), and the neighbouring junctions connected to JOI (Figure 4A, ii), depleting the junctional pool of E-Cad and ensuring that the JOI could recover only from cell-internal stores and not from lateral diffusion from other membranes. Comparing these results to those of single junction FRAP (Figure 4A, i), we observed that *F_s_* was significantly reduced, whereas *F_m_* only showed a small change. In the second approach, we bleached the JOI and large cytoplasmic regions of the cells belonging to JOI, ensuring that internal stores of E-Cad are not available for E-Cad fluorescence recovery (Figure 4A, iii). In these experiments, we observed that the *F_m_* was significantly reduced whereas the *F_s_* only showed a small decrease. Thus, we conclude that lateral diffusion on the membrane contributes to the second-scale recovery fraction (*F_s_)* and this process is mechanical stress independent, whereas membrane trafficking from internal stores contributes to the mechanical stress dependent minute-scale recovery fraction, *F_m_*.

**Figure 4:**
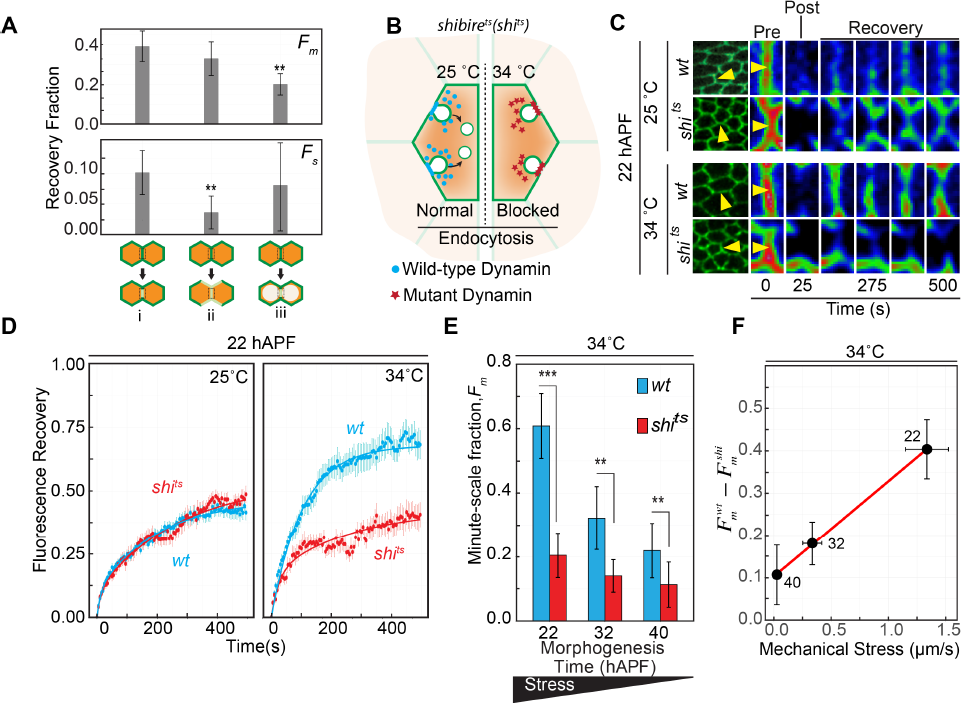
Mechanical stress modulates endocytosis mediated E-Cadherin turnover. **(A)** Minute-scale and second-scale recovery fractions when a single junction is bleached (i), a junction and all its nearest neighbors are bleached (ii), or a junction and the cytoplasm of the contributing cells are bleached (iii). Error bars, SEM. ** represents p < 0.01. P-values are estimated with respect to single junction bleach (i). (n≥ 15 junctions in each case). **(B)** Schematic shows endocytosis block in an temperature sensitive Dynamin mutant, *shi^ts^* upon temperature shift. Wild-type Dynamin shown by blue circles and mutant Dynamin is shown by red stars. **(C)** Time-lapse images from the FRAP time series for wt and *shi^ts^* at 25 ˚C and 34 ˚C shows the pre-bleach, post-bleach and recovery frames. The experiments were performed at 22 hAPF. Yellow arrowheads indicate the junctions that are bleached. **(D)** FRAP curves at 22hAPF for *wt* (blue) and *shi^ts^* (red) imaged at 25˚C (left panel) and 34˚C (right panel). (n≥25 junctions in each case). **(E)** Minute-scale recovery fraction, *F_m_* in *wt* (blue bars) and *shi^ts^* (red bars) at 22, 32 and 40 hAPF. Error bars, S.E.M. ** p < 0.01 and *** p < 0.001. (n≥ 25 junctions in each case). **(F)** Plot of difference in *F_m_* between wt and *shi^ts^* at 34˚C, against mechanical stress, estimated by recoil velocity. Red solid line is the linear fit to the data and r = 0.96 is the Pearson correlation coefficient. Error bars, SEM. See also Figures S10 and S11.

## Mechanical stress modulates endocytosis mediated E-Cadherin turnover

To investigate whether the membrane trafficking fraction (*F_m_)* of E-Cad is derived from endocytosis, we examined the temperature-sensitive Dynamin mutant *shibire^ts^* (*shi^ts^*), in which Dynamin-dependent endocytosis is blocked above the restrictive temperature of 34˚C (van der Bliek and Meyerowrtz, 1991; Classen et al., 2005; De Camilli et al., 1995) (Figure 4B and Figure S10A). Upon shifting *shi^ts^* mutant wings at 22 hAPF, from 25˚C to 34˚C for 10 minutes, mean fluorescence of E-Cad increased at cell junctions suggesting that E-Cad is normally continuously removed from cell junctions by Dynamin-dependent endocytosis (Figure S10B). To ask how blocking Dynamin-dependent endocytosis influenced the minute-scale and second-scale recovery fractions, we performed FRAP on wild-type and *shi^t^s* mutant wings at 22 hAPF, either at 25˚C, or 10 minutes after shifting from 25˚C to at 34˚C (Figures 4C and 4D). Surprisingly, we noted that temperature itself affected E-Cad turnover − in wild-type wings, raising the temperature to 34˚C increases the minute timescale recovery fraction, *F_m_* at the expense of the immobile pool (Figure 4D and Figure S11). In contrast, raising the temperature of *shi^ts^* to 34˚C, decreased *F_m_* and increased the immobile pool (Figure 4D and Figure S11). Thus, endocytic recycling likely contributes to the pool of E-Cad delivered to cell junctions over minute timescales. This idea is supported by our previous observation that both Dynamin and Rab11 are required to maintain a normal distribution of E-Cad at *adherens* junctions specifically at this stage of wing development (Classen et al., 2005). These experiments also suggest that Dynamin dependent endocytosis and recycling of E-Cad increases with temperature.

To investigate whether the endocytosis-dependent pool of E-Cad was affected by mechanical stress, we performed FRAP analysis at 34˚C in wild-type and *shi^ts^* wings at different times of morphogenesis, when mechanical stress gradually decreases (22- 40 hAPF). The difference between *F_m_* in wt and *shi^ts^* at 34˚C, a measure of the contribution of endocytosis to *Fm*, significantly decreased with the decrease in mechanical stress between 22 and 40 hAPF (Figure 4E and Figure S11E). In contrast *F_s_* did not change significantly. Thus, mechanical stress modulates the fraction of E-Cad that turns over through endocytosis (Figure 4F).

## Binding of endocytic regulator p120-Catenin at adherens junctions is controlled by mechanical stress

Since mechanical stress increases endocytic turnover of E-Cadherin, we investigated whether this relation is controlled by the endocytic regulator p120-Catenin (p120). p120-Catenin binds the juxta-membrane domain of E-Cadherin where it interferes with binding of endocytic adaptor proteins and stabilizes E-Cadherin-mediated adhesion (Davis et al., 2003; Myster et al., 2003; Ishiyama et al., 2010; Nanes et al., 2012; Bulgakova and Brown, 2016). We therefore examined whether tissue stress might alter the localization of p120. We used immunostaining to label endogenous p120 (Magie et al., 2002) in the pupal wing epithelium at different stages of morphogenesis. We estimated the junctional enrichment of p120 as the ratio between junctional and cytoplasmic p120 intensities. We found that p120 remains predominantly cytoplasmic when mechanical stress is high and becomes gradually enriched at junctions as mechanical stress decreases (Figures 5A, 5B and 5C). To directly assess the effect of tissue stress on p120 subcellular localization, we compared wild type wings with wings subjected to *dumpy* RNAi. In wild-type wings, when mechanical stress is high at 24 hAPF, p120 is predominantly cytoplasmic. In *dumpy* RNAi wings mechanical stress is reduced at 24 hAPF and the ratio between junctional and cytoplasmic p120 is increased as compared to wild-type (Figures 5D and 5E). This ratio showed a linear correlation with mechanical stress, suggesting that localization of p120 is mechanosensitive (Figure 5F).

**Figure 5:**
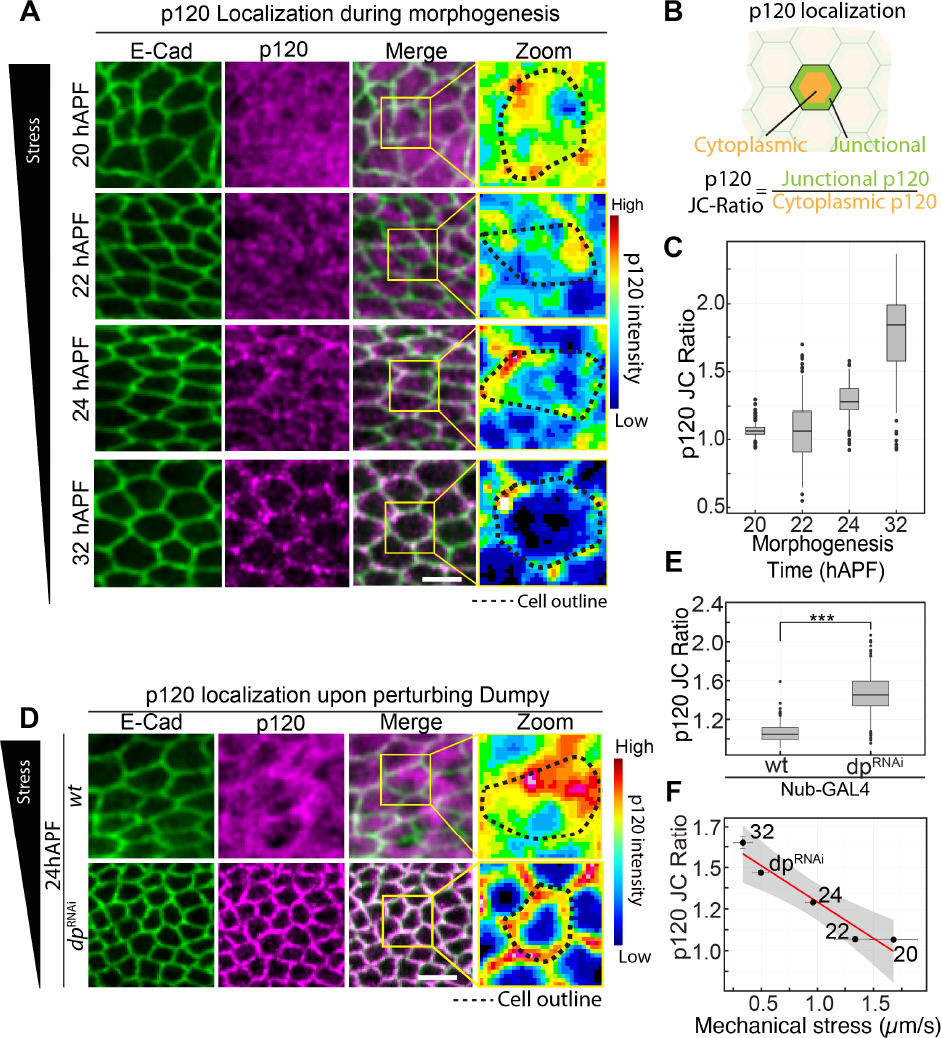
Binding of endocytic regulator p120-Catenin at *adherens* junctions is controlled by mechanical stress. **(A)** E-Cad (green), p120 (magenta) and merge images of the pupal wing at 20, 22, 24 and 32 hAPF stages of wing morphogenesis. The region marked by yellow ROI in the merge image is enlarged to show the color coded p120 intensity. Dotted line indicates the cell outline. Scale bar, 5 µm. **(B)** Schematic shows the estimation of junctional (green) and cytoplasmic (orange) pools of p120 in the cell. The enrichment of p120 in the junctions is estimated as the ratio of p120 intensity in the junctions to the cytoplasm (p120 JC-ratio). **(C)** Box-plot shows the p120 JC-ratio for wt at different stages of pupal wing morphogenesis. (n ≥ 100 cells in each case) **(D)** E-Cad (green) p120 (magenta) and merge images of the pupal wing when wild-type is driven by Nub-GAL4 (top panel) or *dp^RNAi^* is driven by Nub-GAL4 (bottom panel). The region marked by yellow ROI in the merge image is enlarged to show the color coded p120 intensity. Dotted lines indicate the cell outline. Scale bar, 5 µm. **(E)** Box-plot shows the p120 JC-ratio in wt and *dp^RNAi^* wings. *** represent p < 0.001. (n ≥250 cells in each case). **(F)** Mechanical stress (recoil velocity) vs p120 JC-ratio graph for different stages of morphogenesis and dpRNAi. Error bars represent S.E.M.

## p120 regulates mechanosensitive E-cadherin turnover and thereby modifies the viscoelastic behaviour of the wing epithelium

To investigate whether mechanosensitive binding of p120 to E-Cad regulates E-Cad turnover during pupal morphogenesis, we performed FRAP analysis of E-Cad in a p120 null mutant (*p120^308^*) (Myster et al., 2003; Stefanatos et al., 2013). In wild-type wings the minute-scale recovery fraction decreases as mechanical stress decreases during morphogenesis (Figures 6A, 6B and Figure S12). In contrast, in *p120^308^* wings, the minute-scale recovery fraction remains high during morphogenesis (Figures 6A, 6B and Figure S12). This suggests that p120 is required to stabilize E-Cadherin at low mechanical stress thereby making E-Cadherin turnover stress dependent. This raises the possibility that mechanosensitive changes in localization of p120 could mediate the effects of stress on E-Cadherin turnover. To investigate this possibility, we asked whether reduced enrichment of p120 at junctions was responsible for reduced turnover of E-Cadherin in *dumpy* mutant wings. We performed FRAP analysis of E-Cad: GFP in a *p120^308^, dp^ov1^* double mutant background. In *dumpy* mutant wings that are wild-type for p120, the minute-scale fraction of E-Cadherin turnover is lower than in wild-type due to reduced tissue stress (Figures 6C and 6D). Additionally removing p120 from these wings rescues E-Cadherin turnover to wild type levels (Figures 6C and 6D), despite the low mechanical stress in *p120^308^, dp^ov1^* double mutant wings, revealed by laser ablation (Figure S13).

**Figure 6.**
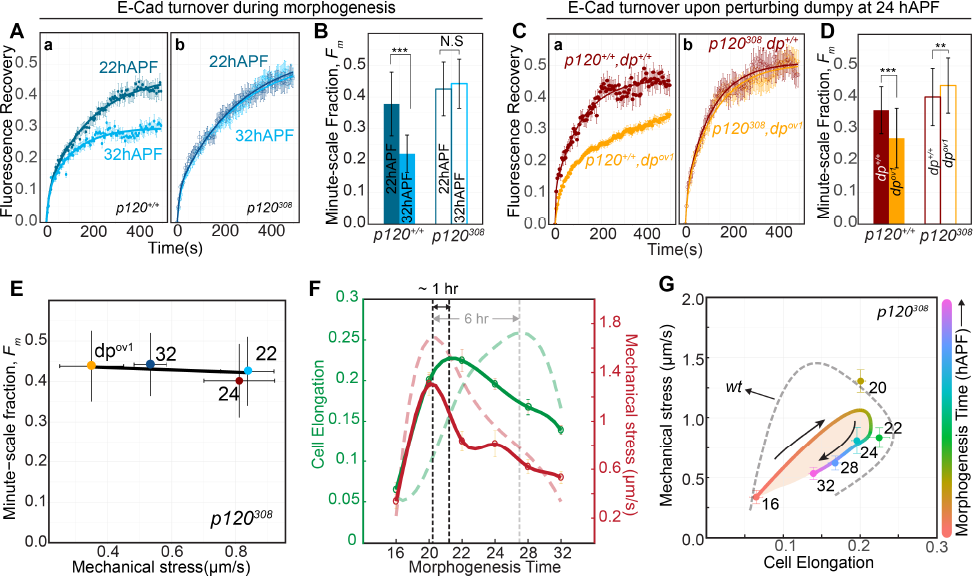
p120 regulates mechanosensitive E-cadherin turnover and thereby modifies the viscoelastic behaviour of the wing epithelium. **(A) (a)** E-cadherin turnover in wild-type (p120^+/+^) at 22hAPF (filled dark blue circles) and 32 hAPF (filled light blue circles). **(b)** E-Cad turnover in *p120^308^* mutant at 22h APF (open dark blue circles) and 32 hAPF (open light blue circles). Solid lines indicate double exponential fits. Errorbars, SEM. (n≥ 30 junctions in each case). **(B)** Minute-scale recovery fraction, *F_m_* for p120^+/+^ at 22 hAPF (Filled dark blue bars) and 32 hAPF (filled light blue bars). *F_m_* for *p120^308^* at 22hAPF(open dark blue bars) and 32 hAPF (open light blue bars). Errorbars, SEM. *** p <0.0001, N.S, not significant. (n≥ 30 junctions in each case) **(C), (a)** E-Cadherin turnover in wild-type (*p120^+/+^,dp^+/+^*) shown in brown and *dp^ov1^* mutant (*p120^+/+^,dp^ov1^*) (orange). **(b)** E-Cad turnover in p120 mutant in wild-type background (*p120^308^,dp^+/+^*) shown in brown open circles, and p120 mutant in dp mutant background (*p120^308^, dp^ov1^*) shown in orange open circles. Solid lines indicate double exponential fits. Error bars, SEM. (n≥ 35 junctions in each case) **(D)** Minute-scale recovery fraction in *p120^308^* mutant in the background of wild-type (brown) and *dp^ov1^* mutant (orange). Error bars, SEM. *** p <0.0001, ** represents p < 0.001. (n≥ 35 junctions in each case) **(E)** minute-scale recovery fraction vs Mechanical stress (recoil velocity) graph for *p120^308^* mutant. (the colored points and the numbers beside them show the different stages of morphogenesis and dumpy perturbation. Solid black line represents the linear fit to the data. Error bars, SEM. **(F)** Changes in cell elongation (green) and Mechanical stress (red) in a *p120^308^* mutant during pupal wing morphogenesis. The open circles represent the data and solid line represents the spline fit through the data. Vertical dotted lines show indicate the time delay of 1 hr between the peak of Mechanical stress and cell elongation. Faint dotted lines show cell elongation (green) and Mechanical stress (red) for wild-type during pupal wing morphogenesis. Faint vertical dotted line indicates the peak of cell elongation in wild-type. Error bars, SEM. **(G)** Mechanical stress vs Cell elongation graph for *p120^308^* mutant. Filled circles are the experimental data color coded for morphogenesis time. The smooth line joining the points is the spline fit to the data color coded for morphogenesis time. Area under the smooth curve (peach) shows the hysteresis in the process and represents the dissipation during the viscoelastic deformation. Faint dotted line shows the hysteresis in Mechanical stress elongation graph for wild-type. See also Figures S12 and S13.

To quantitatively visualize the effect of p120 on stress-dependent E-Cadherin turnover, we plotted the minute timescale fraction of E-Cadherin turnover as a function of mechanical stress, in wings that were mutant or wild-type for p120. We included stress measurements from wt, *dp^ov1^,* and *p120^308^,dp^ov1^* double mutants. In wings, wild-type for p120, there is a strong correlation between mechanical stress and the minute-timescale fraction. This correlation disappears when p120 is lost. (Figure 6E) Thus, these data show that p120 is essential for mechanosensitive E-cadherin turnover.

To explore how p120 might influence tissue viscoelasticity via stress dependent changes in preferred cell shape, we compared cell elongation during the buildup and relaxation of stress in wild type and *p120* mutant wings. In *p120^308^* wings, the delay between the peaks of mechanical stress and cell elongation was only ∼ 1 hour, compared to 6 hours in wild type (Figures 6F and S13). Furthermore, stress-deformation curves enclose a smaller area than wild-type, indicating a more elastic behavior of cells as compared to wild-type (Figure 6G). Thus, these experiments reveal that p120 plays a key role in determining epithelial viscoelasticity.

## Discussion

Epithelial tissues can exhibit distinct responses to applied stresses (Charras and Yap, 2018). In some cases, E-Cadherin mediated adhesions are strengthened in response to stress (Liu et al., 2010), while in other cases they are destabilized (Beco et al., 2015). The molecular mechanisms that underlie stress-dependent strengthening are well studied and involve stress-dependent unfolding of a-Catenin and Vinculin that leads to enhanced linkage to cytoskeleton (Yao et al., 2014; Buckley et al., 2014; Seddiki et al., 2017). However, less was known about the mechanisms that destabilize E-Cadherin. Such stress-dependent changes in E-Cadherin mediated adhesion could affect tissue fluidity or tissue stiffness. Understanding how cells tune the response of E-Cadherin to mechanical stress is therefore important to understand how tissues deform and remodel in response to mechanical stress during morphogenesis. Here, we have shown that the release of p120 Catenin from *adherens* junctions under mechanical stress promotes turnover of E-Cadherin complexes and decreases epithelial viscosity during *Drosophila* pupal wing morphogenesis (Figures 7A and 7B).

**Figure 7:**
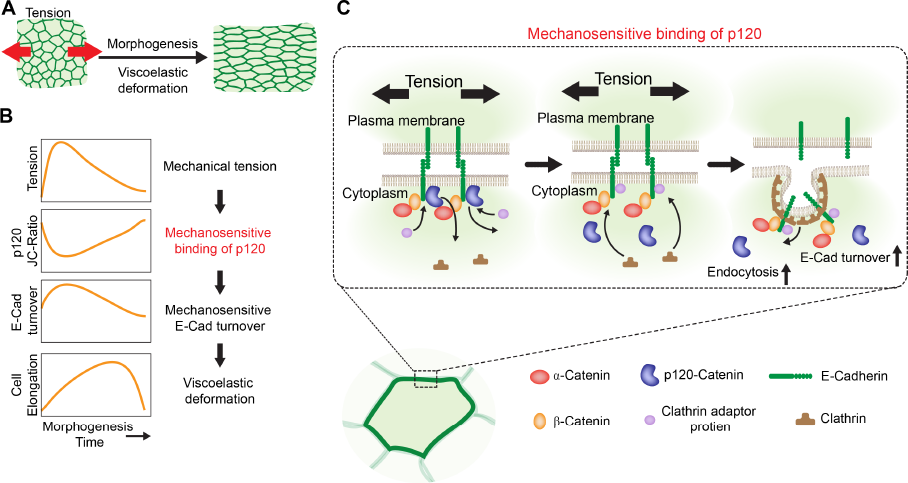
Proposed model for regulation of tissue viscoelasticity by mechanosensitive binding of p120. (A) Viscoelastic deformation of the epithelia during morphogenesis. (B) Steps involved in the viscoelastic deformation. Mechanical stress emerges and decays during morphogenesis. p120 responds to the changes in mechanical stress by changing its junction-to cytoplasmic ratio. This results in mechanical modulation of E-Cadherin endocytosis. This in turn leads to mechanosensitve E-cadherin turnover, essential for viscoelastic deformation of tissues. (C) p120 mechanosensitive binding scheme. Mechanical stress removes p120 from the juxtamembrane domain of E-Cadherin allowing Clathrin adaptor proteins to bind. Binding of Clathrin adaptor proteins recruit Clathrin to the membrane initiating process of endocytosis. Increase in mechanical stress increases p120 mechanotransduction, thereby increasing endocytosis and E-Cadherin turnover.

How does mechanical stress cause p120 to dissociate from *adherens* junctions? One possibility is that mechanical stress-dependent signals produce post-translational modifications of p120 that alter its affinity for the E-Cadherin tail. p120-Catenin can be phosphorylated by a variety of kinases that alter its activity (Mariner et al., 2001; Alemà and Salvatore, 2007; Castaño et al., 2007). Alternatively, mechanical stress-dependent conformational changes in the E-Cadherin cytoplasmic tail might reduce its ability to bind p120 (Figure 7C). If p120 is released by mechanically induced unfolding of the E-Cadherin tail, this would likely require linkage to the actin cytoskeleton via unfolding of a-Catenin and Vinculin. Interestingly, it is precisely these interactions that are required to strengthen E-Cadherin-mediated adhesion in response to stress. This may suggest that stress-dependent internalization could result when forces are exerted for longer times or at higher levels than those required to induce strengthening.

How does release of p120 decrease viscosity? p120-Catenin can bind to the E-Cadherin cytoplasmic tail, and when it does so, it masks binding sites for endocytic machinery (Nanes et al., 2012). Consistent with this, we find that mechanical stress increases the pool of E-Cadherin undergoing endocytic turnover. Faster turnover of E-Cadherin may facilitate cell shape changes and cell rearrangements, reducing epithelial viscosity upon dissociation of p120. p120-Catenin might also impact viscosity through changes in actin/myosin dynamics. When p120 is not bound to E-Cadherin, it is known to modulate the activity of Rho family proteins and the actin cytoskeleton through its interactions with RhoGEF (Noren et al., 2000; Grosheva et al., 2001; Magie et al., 2002). This would be consistent with the observed role for actin turnover in ECadherin stability and the viscosity of cell shape (Engl et al., 2014; Clément et al., 2017).

Our findings show that changes in the level of p120 alter tissue viscosity. In the future it will be interesting to explore whether p120 levels or activity might be used to tune the viscoelastic properties of different tissues during morphogenesis. This might be important in cases where tissues deform at different rates. For example, during *Drosophila* gastrulation, the germband changes its length by 2.5-fold over the course of an hour (Irvine and Wieschaus, 1994), while the *Drosophila* pupal wing requires 18 hours to undergo a smaller deformation. Furthermore, regulating patterns of tissue viscoelasticity could influence the shape that emerges when developing tissues are subjected to mechanical stress.

In conclusion, our findings reveal that mechanosensitive E-Cadherin turnover through p120-Catenin helps determine tissue viscoelasticity. These finding provide new approaches to study how regulation of tissue material properties contributes to tissue shape changes during morphogenesis.

## Acknowledgments

We would like to thank Carl Modes for critical reading of the manuscript. We thank Stephan Grill for providing access to the Laser ablation microscope. This work was supported by the Max Planck Gesellschaft and by a grant from the Deutsche Forschungsgemeinschaft (SPP 1782) to S.E. K.V.I was supported by the ELBE postdoctoral fellowship. R.P.G was supported by ELBE PhD Fellowship. We thank the Light Microscopy Facility of MPI-CBG for help and expert advice.

## Author Contributions

K.V.I, F.J and S.E conceived and designed the project. S.E and F.J supervised the project and participated in data analysis. K.V.I carried out the genetic crosses, FRAP experiments and data analysis, antibody staining and imaging. K.V.I and R.P.G performed laser ablations and data analysis. K.V.I and R.P.G developed the methods and scripts for data analysis. K.V.I, S.E and F.J wrote the manuscript

## Experimental Procedures

### Fly stocks and crosses

The fly strain, *yw; E-Cad::GFP* was a gift from Huang et al (Huang et al., 2009). The stocks of *Nrg::GFP, dp^ov1^, shi^ts1^* were obtained from Bloomington Drosophila Stock Center (stock ID, 6844, 279, 7068 respectively). *UAS::dp^RNAi^* and *UAS::p120^RNAi^* were obtained from Vienna Drosophila Stock Center (Stock ID, v44029). The fly stock *p120^308^* was a gift from Marcos Vidal (Beatson Institute, UK). [*Nub::Gal4, E-Cad::GFP, dp-RNAi(v44029)*], [*shi^ts1^;ECad::GFP*] and [*p120^308^,E-Cad::GFP*], [*p120^308^,dp^ov1^,ECad::GFP]* were generated in the laboratory. Flies were maintained at 25˚C unless specified.

### Laser Ablation experiments

Laser ablation experiments were performed on a Zeiss microscope stand equipped with spinning disk module (CSU-X1; Yokogawa), EMCCD camera (Andor) and a custom built laser ablation system described elsewhere (Mayer et al., 2010). A 355 nm, 1000Hz, pulsed laser was used for ablation.

***Circular ablation*** was performed with a radius of 10 µm in the region between 3^rd^ and 4^th^ longitudinal veins and between the 2^nd^ and 3^rd^ sensory organ from the distal tip of the blade. The laser was focused at equally spaced points (termed as a shot) on the ROI, with a density of 2 shots/µm. For each shot, 25 laser pulses were delivered. Since the wing epithelium has curves and undulations over a area covered by 10 µm radius circle, 5 z-planes, each 1 µm apart were ablated to ensure complete ablation of the circular ROI. In order to capture the recoil of the ablated front along the perimeter of the circular ROI, 5 z-planes, 1 µm apart were acquired with an exposure time of 50 ms and time interval of 670 ms.

***Linear ablation*** was performed in the same region as circular ablation, with a 10 µm line oriented along the antero-posterior axis of the wing, in order to estimate the recoil along the proximal distal axis. The ablation was done only in one single plane as entire region of interest could be acquired in one focal plane. To capture the rapid recoil of the ablated front, single plane images were acquired with 50 ms exposure and the interval between frames was 85 ms. Initial recoil velocity of the ablated region was computed for the estimation of mechanical stress in the tissue.

### Analysis of Laser ablation

***Circular ablation.*** The 5 µm z-stack obtained for each time frame of circular ablation was projected using the maximum projection algorithm of FIJI. An ellipse is fitted to the ablated region, 80 s after ablation and the major and minor axis length and orientation were estimated. Kymographs were generated along the major and minor axis of the ellipse using a FIJI plugin (http://imagej.net/Multi_Kymograph). The initial recoil velocity, which measures the mechanical stress in the tissue, was estimated using a custom written routine in MATLAB (Mathworks). Recoil velocity were measured along the major and minor axis of the ellipse, represented by *v_maj_* and *v_min_* respectively. The orientation of the mechanical stress was estimated from the orientation of the fitted ellipse and the magnitude of recoil velocity along the major and minor axes.

***Linear ablation.***The images obtained from the laser ablations were analyzed using Fiji and MATLAB (Mathworks). Kymographs were drawn on both sides (proximal- and distal-oriented) of the ablated line using the FIJI plugin Multi Kymograph (http://imagej.net/Multi_Kymograph). Both recoil velocities were calculated using a custom written routine in MATLAB (Mathworks). The final recoil velocity (𝑣“#$%”&$) for one ablation was computed as the average of recoil velocities that are both proximal-oriented (*vproximal*) and distal-oriented recoil velocities (*vdistal*):

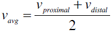

### Particle image velocimetry (PIV)

Junctional displacements after linear laser ablations were analyzed by tracer particles, which in our tissues were GFP-labelled E-cadherin molecules. Flow fields were quantitatively measured using the open-source tool for MATLAB PIVlab (Thielicke and Stamhuis, 2014). The software calculated the displacement of the tracers between pairs of images (before and after the ablation) using the Fast Fourier Transformation algorithm with multiple passes and deforming windows. The interrogation areas were 32 x 32 px^2^ and 16 x 16 px^2^ for the first and second pass, respectively.

### Analysis of cell elongation

Cell elongation in the pupal wing was estimated using the Tissue Analyser plugin in Fiji. The cell elongation for individual cells is calculated as described elsewhere (Aigouy et al., 2010). In order to calculate the cell elongation for the region where laser ablation was performed, vector averaging of the cell elongation of all the cells in the region was performed.

### Fluorescence Recovery After Photobleaching (FRAP)

E-Cad::EGFP line was used to perform FRAP experiments, unless specified. White pupae were collected and aged at 25˚C, unless specified. The pupal case was removed prior to the FRAP experiment as described elsewhere (Classen et al., 2008). While dissecting the pupal case, care was taken to keep the underlying wing untouched and the wing was mounted such that the central region laid flat on the coverslip. FRAP experiments were performed on an Olympus IX81 microscope equipped with a 60x 1.3 NA silicon immersion objective lens (UPLSAPO; Olympus) and a spinning disk scan head (CSU-X1; Yokogawa). Images were acquired on a back illuminated EMMCD camera (iXon DU-897BV; Andor). The region between the second and third sensory organ of the wing from the distal tip, was chosen for FRAP experiments. Multiple single junctions were selected in the field of view and the junctions were bleached using 488 nm laser at 70 % power with dwell time of 50 µs and 2 iterations. Junctions were selected far apart from each other, so that only one junction per cell was bleached. 2-5 pre-bleach and 150 post bleach frames were acquired at 5 sec intervals. Since the pupal wing constantly moves during morphogenesis, a 11 µm z-stack was acquired centered at the bleach plane with 17 planes, 0.7 µm apart. The Z-stack was maximum projected to obtain FRAP movie.

### FRAP combined with laser ablation

FRAP experiments with laser ablation was performed on a Zeiss Axiovert microscope equipped with a Zeiss 63X 1.3 NA water immersion objective and a spinning disc scan head (CSU-X1; Yokogawa). Images were acquired on a back illuminated EMMCD camera (iXon DU-897BV; Andor).

### FRAP Analysis

Since the pupal wing undergoes morphogenesis, the bleached junction in the wing are in constant motion. Before computing the intensity of the junctions after photobleaching, the junctions are corrected for drift. The FIJI plugin “Linear stack alignment with SIFT” (Lowe, 2004) was used to register each frame of the FRAP movie with the previous, to remove the drift form the junctions. Only junctions which show no or very small movement with time were selected for analysis. Background was estimated from the region of the image which does not contain any junctions. Mean intensity of the bleached junctions were estimated over the entire duration of the movie. Reference intensity was estimated by computing the intensity of the entire frame for all frames. Both the junction intensity and reference intensity were background subtracted and normalized by the pre-bleach intensity. The junction intensity normalized by the reference intensity to obtain the intensity corrected for bleaching during acquisition was computed as,

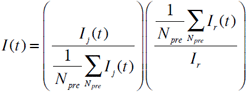

where *I_j_(t)* and *I_r_(t)* represent the junction intensity and reference intensity respectively and *N_pre_* is the number of pre-bleach frames.

This was further normalized as:

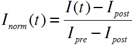

where

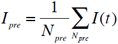

## Bootstrapping parameter estimation

Bootstrapping procedure was used to fit the FRAP curves and estimate the recovery parameter. Towards this, 3 FRAP curves were randomly selected from the set of FRAP curves for a given experiment and their mean was estimated. A double exponential of the following form was fitted to the curve and the fitting parameters were estimated:

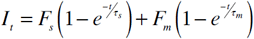

where *I_t_* is the normalized fluorescence intensity at time t, *F_s_* is the second-scale recovery fraction, *F_m_* is the minute-scale recovery fraction, t_s_ is the second-scale timescale and t_m_ is the minute-scale timescale. A double exponential provided the best fit to the curve (Table S1). This procedure was performed iteratively for 1000-2500 iterations to obtain a smooth distribution of the fit parameter.

## Temperature Shifts

Temperature shifts were performed to activate the temperature sensitive mutant, *shi^ts1^*, referred as shi^ts^. Both *shi^ts^* and *wt* flies were raised in 25˚C. White pupae were collected and were aged till the right pupal stage at 25˚C. The mutant *shi^ts^* is inactive at the permissive temperature of 25˚C, whereas its active at restrictive temperature of 34˚C. So, FRAP experiments were performed in both *wt* and *shi^ts^* at 25˚C (control) and 34˚C (experiment). Control pupae were maintained at 25˚C throughout the FRAP experiment. For experiment, pupae were shifted from 25˚C to 34˚C at the appropriate stage of pupal development, on the microscope and incubated for 10 min, to activate the *shi^ts^* mutation. This was followed by the FRAP experiment in either *wt* or in *shi^ts^*.

## Immunofluorescence staining of p120

E-Cad::GFP white pupae were aged at 25˚C to appropriate time point and then the pupal wing dissected in PBS as described elsewhere (Classen et al., 2008). The dissected wings were fixed in 8% Paraformaldehyde (PFA) for 30 min. The wings were then washed twice in PBT (PBS + 0.04%Triton X-100) for 5 min each. This was followed by incubating the wings in PBT2 (PBS + 0.2% Triton X-100) for 20 min. Then the wings were blocked in PBTN (PBT + 4% NGS) for 20 min before they were incubated in anti-*p120* antibody prepared in PBTN (1:2 dilution) overnight at 4˚C. The anti-p120 Catenin antibody (p4B2) was deposited to DSHB by Parkhurst, S. (DSHB, Hybridoma Product p4B2) (Magie et al., 2002). Then the wings were washed in PBT 3 times quickly followed by 3 times for 10-15 min. Then wings were incubated in goat anti-mouse Alexa-647 secondary antibody (Life technologies) prepared in PBTN (1:500 dilution) for 4 hours in a rocking platform at room temperature. The secondary antibody was removed and the wings were washed in PBT 3 times quickly followed by 3 times for 10-15 min. Then the wings were washed once in PBS. The wings were then sucked up with a 100 µl tip and transferred onto a glass slide. The wings were then mounted using Vectashield (Vector Laboratories) mounting medium and stored at 4˚C.

## Imaging and analysis of p120 localization

An Olympus IX83 microscope equipped with Yokogawa CSU-X1 spinning disk module was used to image the p120 immunofluorescence samples. The samples were illuminated using 488 nm laser for imaging EGFP and 638 nm laser for imaging Alexa-647. Images were acquired using 525/50 (for EGFP) and 685/40 (for Alexa-647) bandpass filters on an Andor EMCCD camera (iXon 888). Localization of p120 within cells was estimated using Fiji plugins and custom written MATLAB routines. The junctional network visualized by E-Cadherin::GFP was segmented using Tissue Analyser. A 3 pixel junctional mask (*M_j_*) was generated for each cell and was multiplied with the p120 junctional intensity of p120 (*p120_junc_*). A mask of the cell cytoplasm (*M_c_*) was generated by subtracting the mask for the junctional network from the whole cell mask. This *M_c_* was multiplied with the p120 image to get the cytoplasmic intensity of p120 (*p120_cyto_*). The enrichment of p120 in junction was given by the p120 junction to cytoplasm ratio (p120 JC-Ratio) as

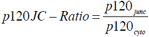

### Statistical tests

Student’s t-test was performed to estimate significance of the quantities observed. The corresponding p-values are mentioned in the figure legends.

